# The influence of eye movements and their retinal consequences on bistable motion perception

**DOI:** 10.1101/2020.08.07.241091

**Authors:** Mareike Brych, Supriya Murali, Barbara Händel

## Abstract

Eye related movements such as blinks and microsaccades are modulated during bistable perceptual tasks, however, the role of such movements in these purely internal perceptual switches is not known. We conducted two experiments involving an ambiguous plaid stimulus, wherein participants had to continuously report their motion percept. To dissociate the effect of blinks and microsaccades from the visual consequences of such eye movements, we added external blanks and microshifts.

Our results showed that while blanks facilitated a switch to the coherent motion percept, this was not the case for a switch to component percept. A similar difference was found with respect to blinks. While both types of perceptual switches were preceded by a decrease in blinks, only the switch to coherent percept was followed by an increase in blinks. These blink related findings, which we largely replicated and refined in a second study, indicate distinct internal processes underlying the two perceptual switches. Microsaccade rates, on the other hand, only showed a weak relation with perceptual switches but their direction was modulated by the perceived motion direction. Additionally, our data showed that microsaccades are differently modulated around internal (blinks) and external events (blanks, microshifts), indicating an interaction between different eye related movements.

This study shows that eye movements such as blinks and microsaccades are modulated by purely internal perceptual events independent of task related motor or attentional demands. Eye movements therefore can uncover distinct internal perceptual processes that might otherwise be hard to dissociate.

## Introduction

Although eye movements like microsaccades and blinks are known to be influenced by sensory input (Bonneh et al., 2016), it is not clear how they are related to purely internal perceptual processes. One way to study this is by looking at ambiguous stimuli. Ambiguous or bistable percepts occur when a single unchanging stimulus can be interpreted in two or more ways. The most well-known bistable stimuli are the Necker cube (Necker, 1832) and the face-vase illusion (Rubin, 1921)., Also, moving stimuli exist, which can be perceived as moving in different directions, e.g. ambiguous motion quartet (von Schiller, 1933) and the ambiguous plaid stimulus (Adelson & Movshon 1982). The different interpretations of the ambiguous stimuli are usually exclusive and alternating, though mixed percepts do occur.

Although, a modulation of eye movements during the perceptual switches has been observed before (Einhauser et al., 2008; Ito et al., 2003; Laubrock et al., 2008; Nakatani et al., 2011; Nakatani & van Leeuwen, 2005; Troncoso et al., 2008; van Dam & van Ee, 2005, 2006), it is still not clear if these eye movements are a consequence of or play a causal role in the switch. Ito et al. (2003) for instance, found that no unique eye behavior was associated with switching, but that saccades and blinks were sensitive to either the perceptual change or the response preparation respectively. Specifically, they found that saccades increased before the response, whereas blinks decreased before and increased after the response. The reduction in blinks before and increase in blinks after the response has been replicated by others (van Dam & van Ee, 2005, 2006). In fact, van Dam and van Ee (2005) argue that blinks are synchronized to all task relevant events and that the switch and the response to the switch might act as such events.

It must be noted that only a few studies analyzed the modulation of blinks in a time resolved manner and used a reasonably large sample size (Ito et al., 2003). Other studies that did conduct a time-resolved analysis did not look at blinks, but rather focused on saccades and microsaccades (Einhauser et al., 2008; Laubrock et al., 2008; Nakatani et al., 2011; Otero-Millan et al., 2012; Troncoso et al., 2008). Our aim was to look at blinks and microsaccades in a time-resolved manner and extend previous findings to a larger sample size.

Microsaccades do play a role in perception as they have e.g. been shown to prevent visual fading that can occur during fixation (Martinez-Conde et al., 2006). With regard to the role of microsaccades during bistable perception, some studies have found a pattern similar to blinks, with a reduction before the response iniacting a perceptual switch, most likely during the switch itself, and an increase after the response (Laubrock et al., 2008; van Dam & van Ee, 2006). However, others have suggested a more causal role in the perceptual transitions with an increase and not a decrease before the switch. For instance, through a time-resolved analysis, Otero-Millan et al. (2012) found that microsaccade increase before perceptual switches in the rotating snake illusion and Troncoso et al. (2008) found that faster motion perception while viewing the Enigma illusion was associated with higher microsaccade rates.

The role of microsaccades (and larger saccades) has also been observed in terms of their direction, specifically during bistable motion stimuli. In general, it has been observed that saccade directions coincide with the perceived direction of motion (Baker & Graf, 2010; Laubrock et al., 2008). In fact, Baker and Graf (2010) found that asking subjects to make eye movements congruent with a specific motion percept, prolonged that percept.

Our goal was to understand if eye movements are a consequence of perceptual transitions or if they rather cause these transitions. We further compared the effect of eye movements to the influence of their retinal consequences. Specifically focusing on blinks and microsaccades, the retinal consequence of a blink is a transient interruption of the visual scene, and the consequence of the microsaccade is a shift in the image. By introducing blanks and microshifts (see Methods), we could dissociate the effect of the eye movement itself from their subsequent consequence on the visual scene. Although multi-second interruptions in ambiguous stimuli has been shown to stabilize percept (Leopold et al., 2002; Noest et al., 2007), interruptions that mimic a blink in terms of timing and rate have not been investigated.

In two experiments we used the ambiguous plaid stimulus, which consists of moving gratings, superimposed over each other (Hupé & Rubin, 2004; Wallach, 1935). The stimulus is seen either as one single grating (coherent percept) or as two separate gratings (component percept). The ambiguity arises due to the aperture problem: a one-dimensional spatial structure cannot be determined unambiguously if it is viewed through a small aperture such that the ends of the stimulus are not visible (Binder et al., 2008).

Our results showed that eye movements and their visual consequences have different influences on perceptual switches and that the modulation of eye blinks is mediated by attention.

## Materials and methods

The study consisted of two experiments using a very similar stimulus and setup. Experiment 2 was conducted as a replication study and additionally used an eye tracker that allowed us to analyze microsaccades in addition to blinks.

### Stimulus

A moving grating displayed behind a fixed-size aperture is usually perceived as moving in a perpendicular direction. Superimposing another grating with a different orientation creates the ambiguous plaid stimulus: The two gratings can either be perceived on top of each other as two components with different directions or as a unified plaid pattern coherently moving in one direction. For the stimulus with 0° rotation, the coherent plaid pattern moved directly downward, the components moved in angles of ±67.5° to the coherent motion direction (Fig. 1A). The size, speed and rotation of the moving grating is specified for each experiment, separately (please see below).

**Figure 1.**
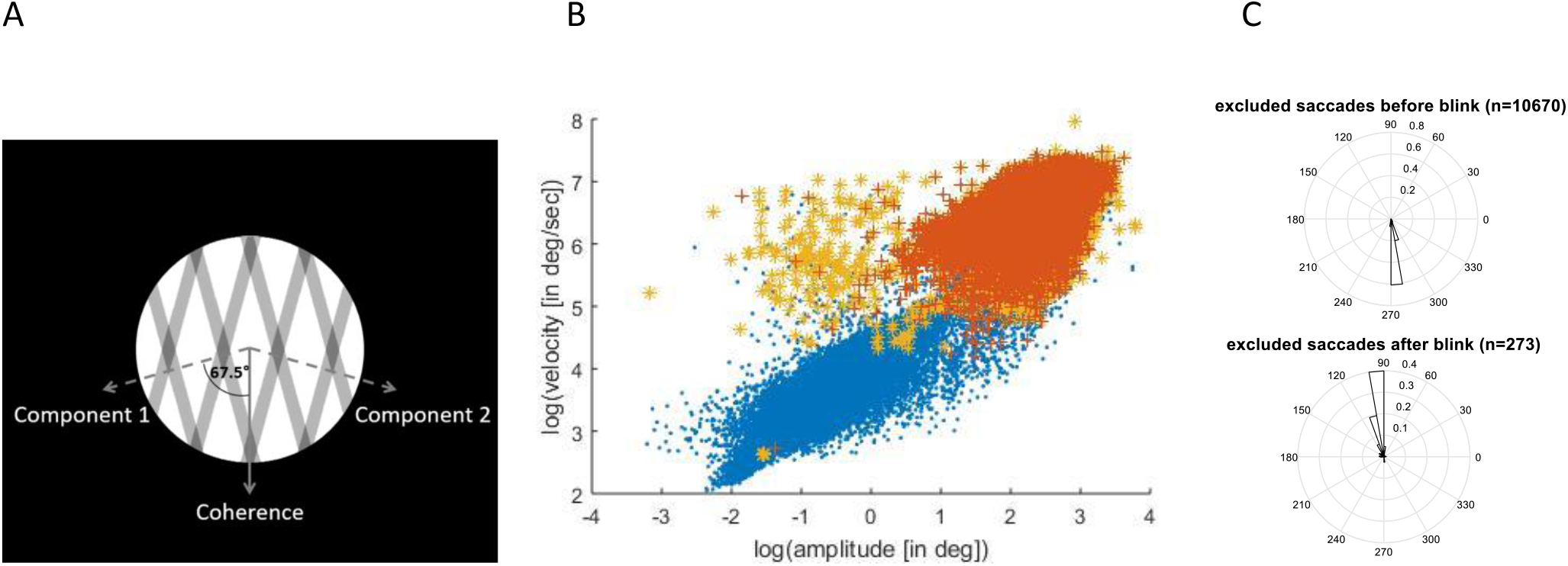
A. Stimulus representation. B. Saccade detection. Yellow stars mark saccades, which were excluded based on the velocity criterion. Red crosses mark saccades, which were detected ± 20 ms around a blink. Blue dots represent valid (micro-)saccades. C. Direction of detected saccades 20ms before or after a blink

The experimental program was implemented in MATLAB, using the Psychophysics Toolbox extensions (Brainard, 1997; Kleiner et al., 2007; Pelli, 1997). Response buttons were stuck on the table being connected to a BBTK response box (model: K-RB1-4; The Black Box ToolKit Ltd, UK), which was connected to a laptop via USB. Participants indicated their prevalent percept by continuously pressing one of two buttons with their right index finger only. Lifting the button was the first indicator that the percept changed, which is why we present our results in relation to the button lift instead of the later happening button press.

### Experiment 1

#### Participants

Fourteen psychology students of the University of Würzburg (age: 20.5 ± 2.18 years (mean ± SD)) took part in the first experiment. They received study credit for their participation. All participants had normal or corrected-to-normal vision. The study was approved by the local ethics committee and complied with the European data protection law (DSGVO). The participants gave their written informed consent before taking part in the study.

#### Procedure

Participants were seated in a dark room 40cm away from the screen with their heads fixed using a chin rest. For stimulus presentation, we used a NEC MultiSync monitor (1024×768 resolution, 60 Hz refresh rate), which was controlled by a Dell Precision (M6700) laptop running Windows 10. Binocular eye movements were recorded with 120 Hz using SMI eye tracking glasses (SensoMotoric Instruments GmbH, Berlin, Germany).

The stimulus was presented in an aperture of 10.0° in diameter, on a black background and moved with a speed of 1.3°/ sec. A red fixation spot of 0.43° in diameter was placed in the center of the stimulus (Fig. 1 A).

During a spontaneous eyeblink, the eyeball has been reported to slightly move downward and inward (Collewijn et al., 1985). To test for a possible influence of the vertical movement of the eyeball during blinks on the percept, four different stimulus rotations were presented: 0°, 67.5°, 112.5° and 180°, with respect to the coherent motion. These specific rotations were chosen such that either the coherent motion of the gratings moved up (180°) or downwards (0°), or one of the gratings (components) moved up (112.5°) or downward (67.5°).

To control for the visual changes during a blink, the screen was blackened for a random duration between 116ms and 167ms randomly every three to six seconds (in steps of 0.5s) in half of the trials, which will be referred to as “blanks”.

The first experiment consisted of eight trials with a duration of six minutes each. Each of the four stimulus rotations were presented twice, once with blanks once without. Trial order was completely randomized. A 1-point calibration of the SMI eyetracker was performed prior to the start of the experiment.

### Experiment 2

#### Participants

Twenty-two new participants (15 females, age: 27.4 ± 8.88 years (mean ± SD)) took part in the second experiment. They received payment or study credit for their participation. All participants had normal or corrected-to-normal vision. They gave their written informed consent prior to the participation. The study was approved by the local ethics committee and complied with the European data protection law (DSGVO).

#### Procedure

Participants were seated in a darkened room and placed their head on a chin rest 68cm away from the monitor. The stimulus was presented on a Mitsubishi Diamond Pro 2070SB monitor (1152×864 resolution, 60 Hz refresh rate). The experiment was controlled by a Tuxedo laptop running Ubuntu 16.04 LTS. To analyze very small eye movements, binocular eye movements were recorded with 500 Hz using an EyeLink 1000 eyetracker (SR Research, Ottawa, Ontario, Canada).

The stimulus had a diameter of 5.8° and moved with a speed of 0.7°/ sec. The fixation spot was 0.25° in diameter.

Again the influence of the vertical movement of the eyeball during a blink and the horizontal movement of the eyeball during a microsaccade were controlled with four stimulus rotations. We used again the stimulus rotations of 0° & 67.5° (blink related) and added rotations of 22.5° (one component moving horizontally to the right, the other one to the bottom right) and 90° (coherent motion horizontally to the right) (microsaccade related).

Additionally, to the blanking trials, microshift trials were introduced to control for the visual changes during a microsaccade. During microshift trials, the stimulus randomly shifted every three to six seconds randomly towards the right or the left by 0.2°. The fixation spot stayed at its position. The maximal deviation of the stimulus from the original position was 0.8°, i.e. the shift could maximally happen 4 times in the same direction.

The second experiment consisted of two blocks each having eight trials of three minute duration in random order. Before each block, a 5-point calibration and validation of the Eyelink eye tracker was performed. Each block consisted of four test trials (all four rotations described above), two blank trials to simulate blinks (rotations 0° and 67.5°) and two microshift trials to simulate microsaccades (rotations 22.5° and 90°).

### Data analyses (Experiment 1 and 2)

For the first experiment, two participants were excluded because of a lower blink rate than 5 blinks/min, another one due to more than 39% of missing data. For the second experiment, one participant was excluded due to very high blink rate (>35 blinks/min) and two more due to a blink rate lower than 5 blinks/min during all trials without blanks or microshifts.

The identical trials of block one and two in the second experiment were concatenated before analysis. Events (blinks, blanks, microsaccades, microshifts) onsets were counted for bins every 100ms around the button lifts indicating perceptual switches. If such an event was detected around multiple switches, we divided the counts by the number of occurrences. To incorporate different switch rates, we averaged the time course over all switches. Furthermore, we controlled for different rates of eye movements by dividing the result by the number of counted eye movements. Finally, these time series were z-transformed. These time series around a switch were compared to time series were no switch occurred.

Cohen’s d for paired samples t-tests was calculated as the mean of D divided by the standard deviation of D, where D is the differences of the paired samples values.

For statistical analysis, we implemented the nonparametric statistical test described by Maris and Oostenveld (2007) which is based on clustering of adjacent time samples that show a similar difference in sign and magnitude. The threshold was selected as the 97.5 quantile of a T-distribution.

### Blink detection (Experiment 1 and 2)

Whenever the eyelid covers the eye, rapid changes in pupil diameter are recorded by the eyetracker. Therefore, we developed a blink detection algorithm based on pupil diameter. Firstly, pupil diameter was z-transformed and mean and standard deviations were computed. By visual inspection, a manually set amount of standard deviations (between 1.9 and 4) was chosen for threshold. A blink was assumed when z-transformed pupil diameters of both eyes decreased below this threshold or if the pupil was not detected at all. The start and the end of the blink were then extended until the diameter of both eyes were higher than half the threshold. Blinks less than 100ms apart from one another were concatenated and blinks shorter than 50ms or longer than 1000ms were discarded.

### Microsaccade detection (Experiment 2)

We implemented an algorithm based on the description by Engbert and Kliegl (2003) where a transformation of fixation positions to two-dimensional velocity space is performed to detect microsaccades according to their dynamical overshoot component. We assumed a minimal duration of four samples (8ms) and only considered binocular microsaccades up to an amplitude of 1°. In line with previous research, (micro-)saccades showed a linear relation of amplitude and peak velocity known as “main sequence” (Zuber et al., 1965). Furthermore, we excluded (micro-)saccades which had a velocity more than three absolute deviations away from the median (Leys et al., 2013) (Fig. 1B, yellow stars). The velocity criterion excluded 0.2% of all microsaccades. As shown in figure 1B, these criterions (amplitude and velocity based) excluded also most of those blink-induced events that might have been mistaken for a saccade. When plotting events detected as saccades which happened within 20 ms of a blink (Fig. 1B, red crosses) we see that they form a separate cloud showing a much larger amplitude with higher peak velocity compared to the otherwise prevalent microsaccades during fixation (Fig. 1B, blue dots). The interpretation, that these detected events indeed were not genuine saccades but rather blink-induced, is further strengthened by the finding that they primarily show a downward direction when happening before the blink and an upward direction when happening afterwards (Fig. 1C). This stands in gross contrast to the preferred horizontal direction of microsaccades (Laubrock et al., 2005; Valsecchi et al., 2007). Only 0.01% of saccades after exclusion would fall within 20 ms of a blink and were excluded additionally (as in (McCamy et al., 2013).

## Results

During the first experiment, participants perceived coherent motion longer than component motion (21.03 ± 21.83 sec compared to 8.06 ± 8.03 sec (mean ± standard deviation); paired t-test: *t*(10)=2.16, *p* = .056, *d* = 0.65). Coherent motion was also dominant in the second experiment (*t*(18) =4.61, *p* < .001, *d* = 1.06), but both percept durations were slightly shorter (14.77 ± 6.65 sec compared to 7.57 ± 4.41 sec (mean ± SD)). The duration of percept was calculated between a button press and the corresponding lift and revealed the typical unimodal and positively skewed distribution when plotted as histograms (not shown).

### Blink/blank results (Experiment 1 and 2)

During the first experiment, participants blinks on average 11.12 ± 5.39 (SD) times per minute with a mean duration of 136.80 ± 26.80 ms (SD). For the second experiment, the blink rate was 12.09 ± 8.83 (SD) blinks per minute with a mean blink duration of 171.32 ± 46.47 ms (SD). Furthermore, we calculated the blink rate separately for the different percepts taking into account the respective percept durations. During both experiments, participants blinked significantly more during coherent motion than during component motion (experiment1: *t*(10) = 5.56, *p* < .001, *d* = 1.68; experiment 2: *t*(18) = 4.86, *p* < .001, *d* = 1.12). Blanks were slightly shorter than blinks in both experiments with a mean length of 125 ± 19 ms (SD) and 141 ± 19 ms (SD) respectively.

**Figure 2.**
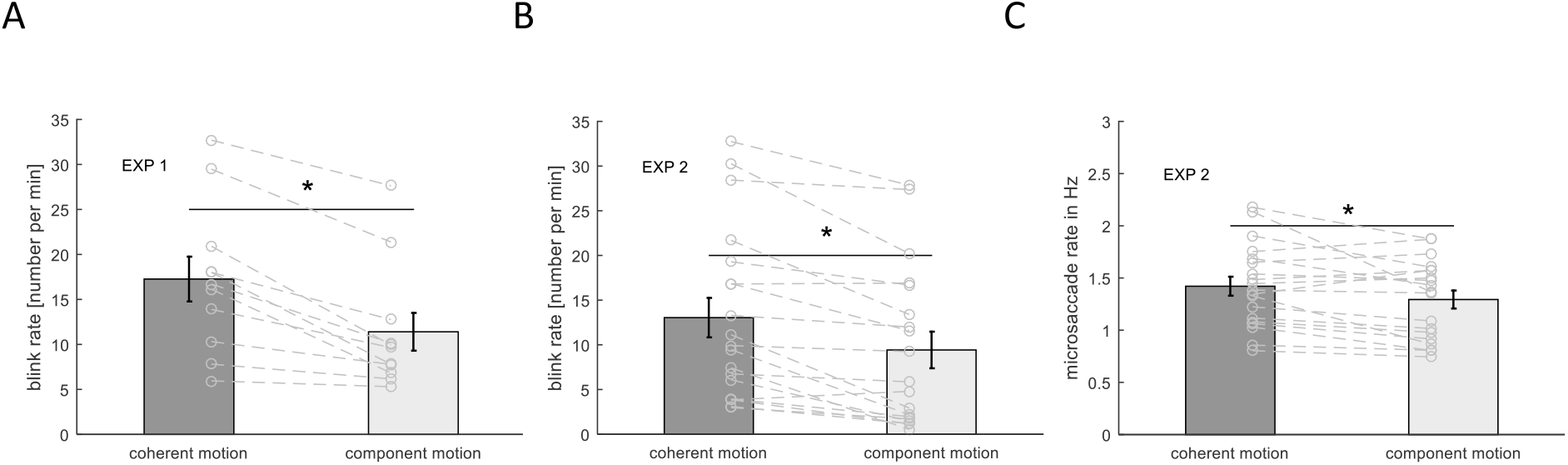
A. Blink rate during the different percepts in experiment 1. B. Blink rate for the different percepts in experiment 2. C. Microsaccade rate for the different percepts in experiment 2. Bars and error bars represent mean ± standard error of the mean (SEM). Gray lines represent data of individual participants. An asterisk marks a significant difference at *p* < .001.

To investigate if a change in perception is linked to a blink event, we looked at the normalized blink rate around perceptual switches, separately for switches to coherent and component motion and statistically compared it to the normalized blink rate when no switch occurred. The same was done for the normalized blank rate. This was done to assess if any influence was introduced by the visual consequences of the eye closure during a blink, as mimicked by the blank. Normalized blink rates were taken from trials without blanks or microshifts, but were combined over stimulus rotations. Please note that no p-values are reported due to the non-parametrical statistical testing that was applied (Maris and Oostenveld (2007).

When switching to coherent motion, there was a significant decrease in blink rate between −800 and −200 ms before the button lift (indicating a perceptual switch) in the first experiment compared to time periods with no switch. This decrease was replicated in the second experiment, where we found significant differences between −700 and −400 ms (Fig. 3). When switching to component motion, such a decrease in blink rate was found in the first experiment (−300 to 0 ms before the switch), but did not reach significance in the second experiment, although a decrease before the switch is clearly visible. Interestingly, blink rate strongly increased around the time of a response indicating a switch to coherent motion, possibly at the time of the perceptual switch. This peak in blink rate is clearly visible in both experiments, but statistical comparison between blink rate around a switch and around no switch only reached significance in the second experiment between 300 and 600 ms after the response.

**Figure 3.**
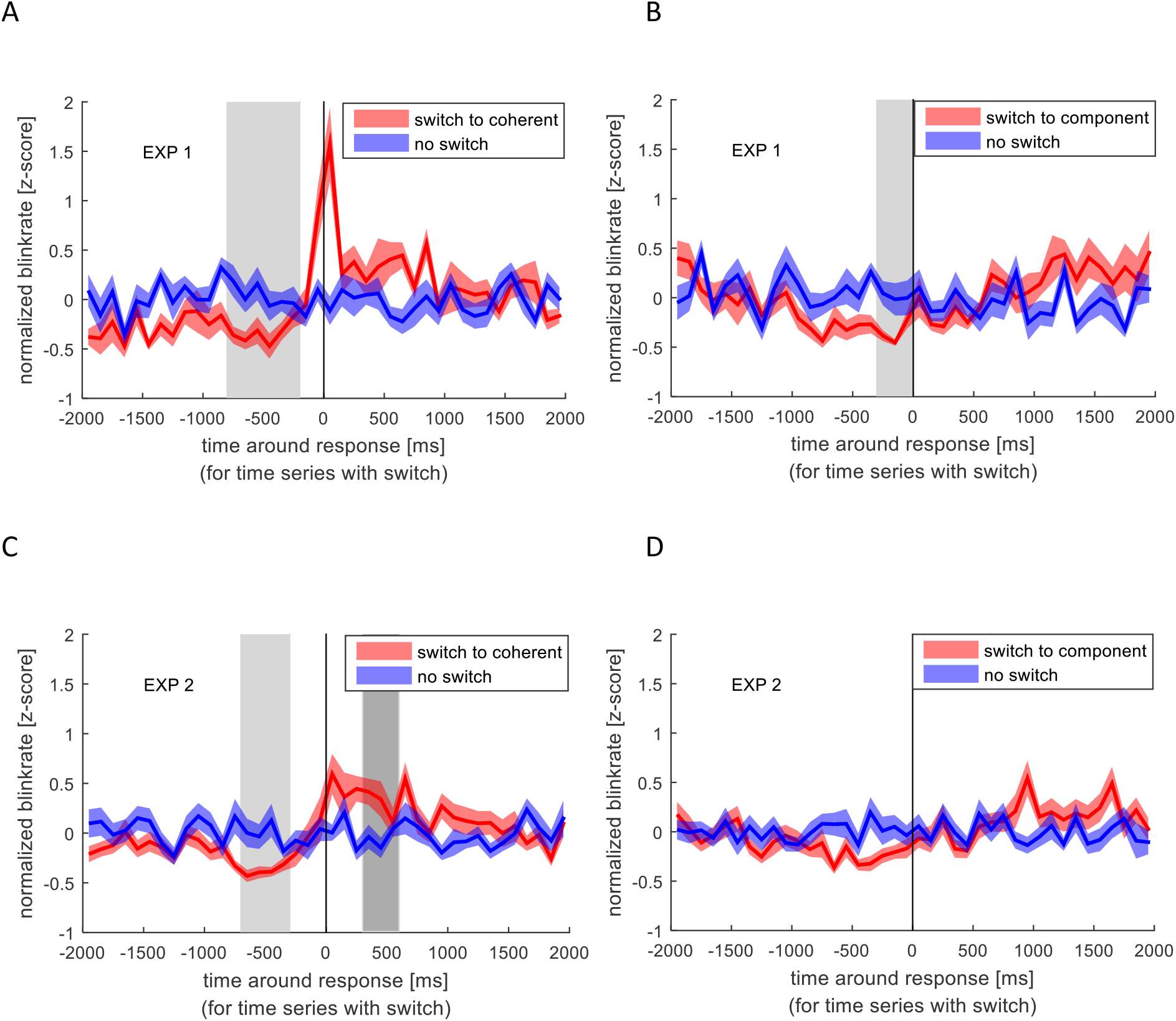
Normalized blink rate around the response indicating a perceptual switch (red) compared to the normalized blink rate during no switch (blue). Colored lines and ribbons represent mean ± standard error of the mean (SEM). Vertical shaded areas mark significant time points revealed by the non-parametrical statistical test procedure described by Maris & Oostenveld (2007). A. Switch to coherent motion in experiment 1. B. Switch to component motion in experiment 1. C. Switch to coherent motion in experiment 2. D. Switch to component motion in experiment 2.

In contrast to the blink rate modulation, blanks, albeit again showing the strongest effect for switches to coherent motion, showed a different temporal pattern. As shown in Figure 4, the blank rate increased before the switch to coherent motion in experiment 1, which was even more pronounced in experiment 2. This increase in blank rate around the switch to coherent motion was significantly different from the blank rate around no switch between −900 and −500 ms before the response in the second experiment. The blank increase preceded the perceptual switch and was not predictable. This pattern was not visible when switching to component motion. When looking at the different stimulus rotations separately, all patterns were very similar which means that the effect of blanks and blinks are independent of the movement direction of the stimulus.

**Figure 4.**
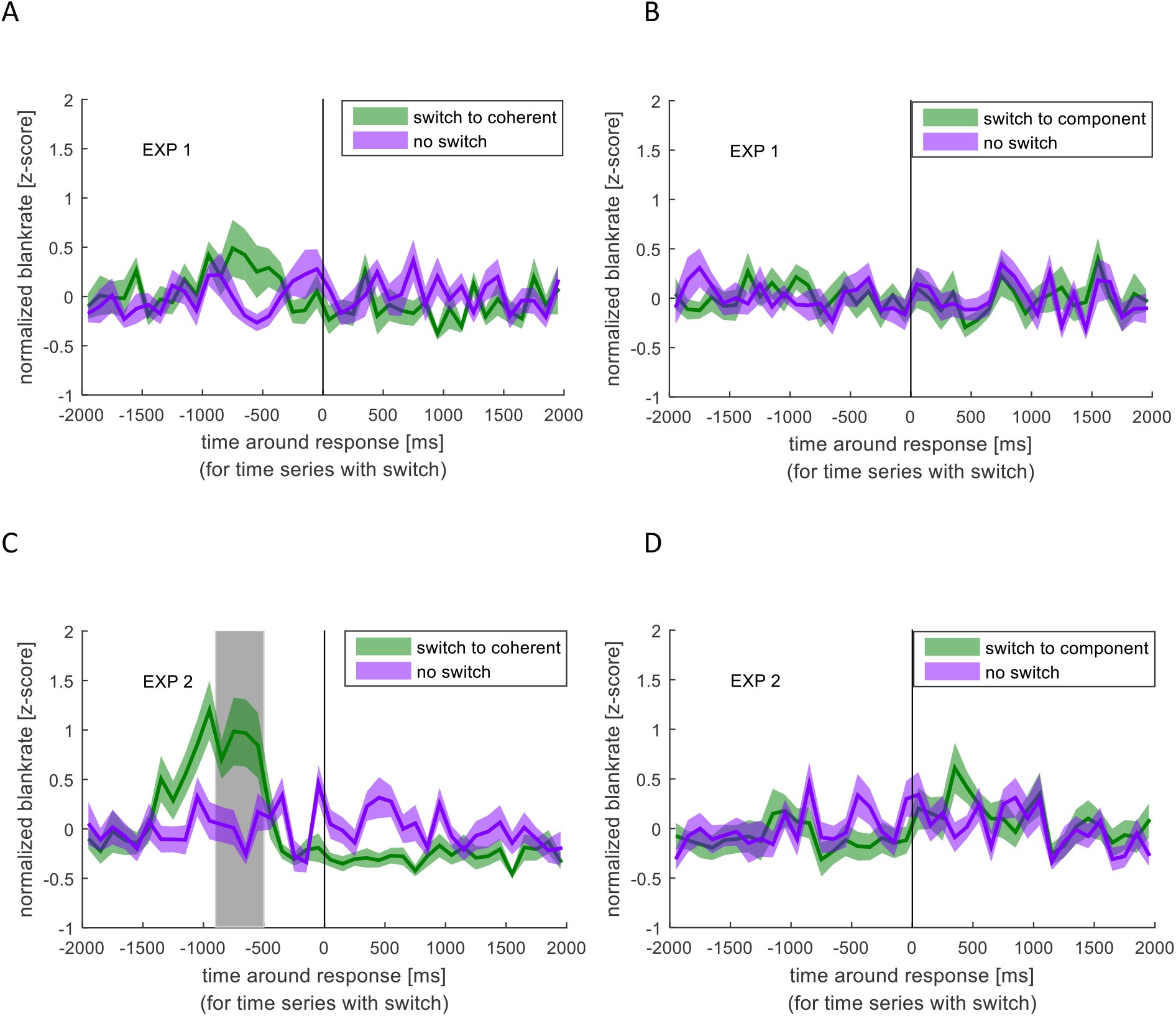
Normalized blank rate around the response indicating a perceptual switch (green) compared to the normalized blank rate during no switch (purple). Colored lines and ribbons represent mean ± standard error of the mean (SEM). Vertical shaded area marks significant time points revealed by the non-parametrical statistical test procedure described by Maris & Oostenveld (2007). A. Switch to coherent motion in experiment 1. B. Switch to component motion in experiment 1. C. Switch to coherent motion in experiment 2. D. Switch to component motion in experiment 2.

### Microsaccades (Experiment 2)

Due to the low sampling frequency of the eyetracker used in experiment 1, we were only able to analyze microsaccades in the second experiment.

Over all trials and participants, we found a microsaccade rate of 1.39 ± 0.40Hz (mean ± SD). Looking at the different percepts, a paired t-test revealed that participants had a significantly higher microsaccade rate during coherent motion than during component motion (*t*(18) = 2.43, *p* = .026, *d* = 0.57) taking into account the respective percept durations (Fig. 1C). Coherent percept is therefore associated with a higher microsaccade rate as well as a higher blink rate as compared to the component percept.

Similar to the comparison of normalized blink rate around perceptual switches and no switches, we looked at the differences between normalized microsaccade rate around switches and no switches. To assess the specific influence of the visual shift introduced by a microsaccade (as mimicked by the microshift), we compared microshift rate around switches and no switches (Fig. 5).

**Figure 5.**
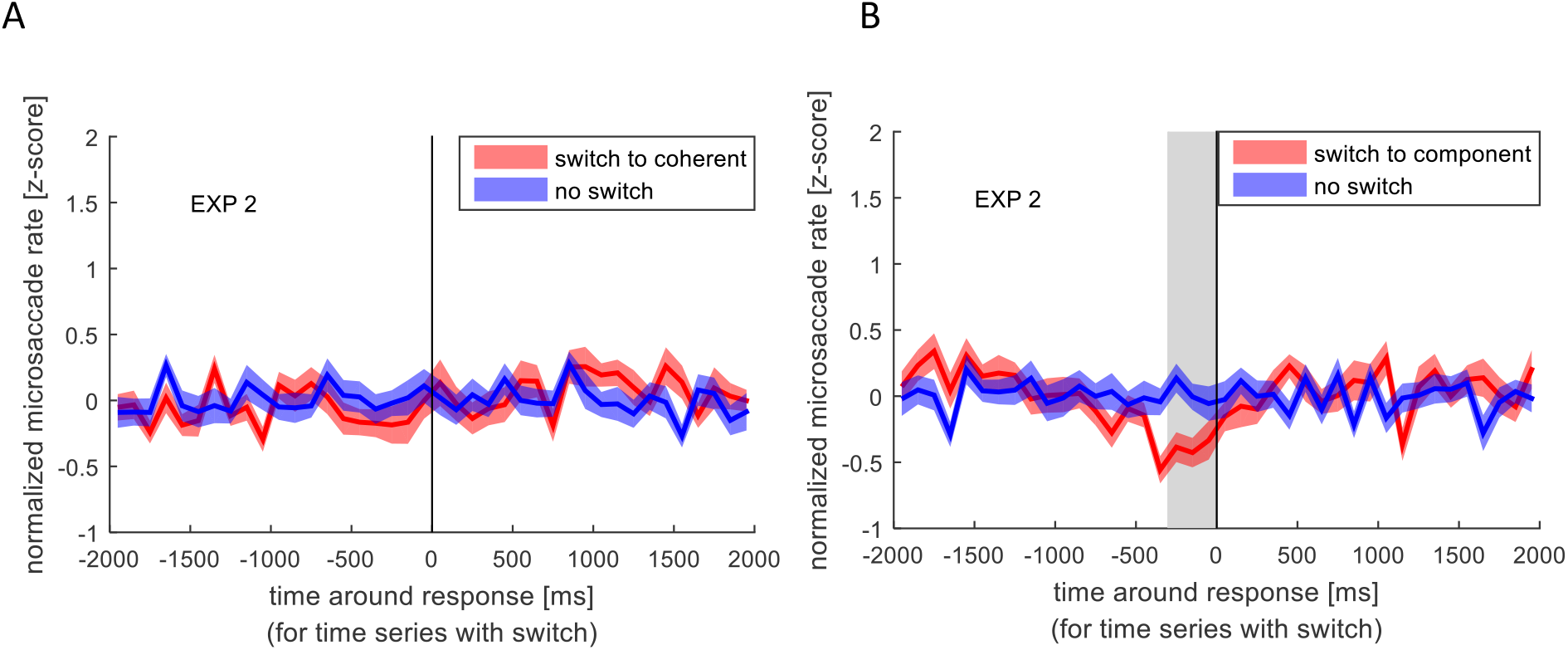
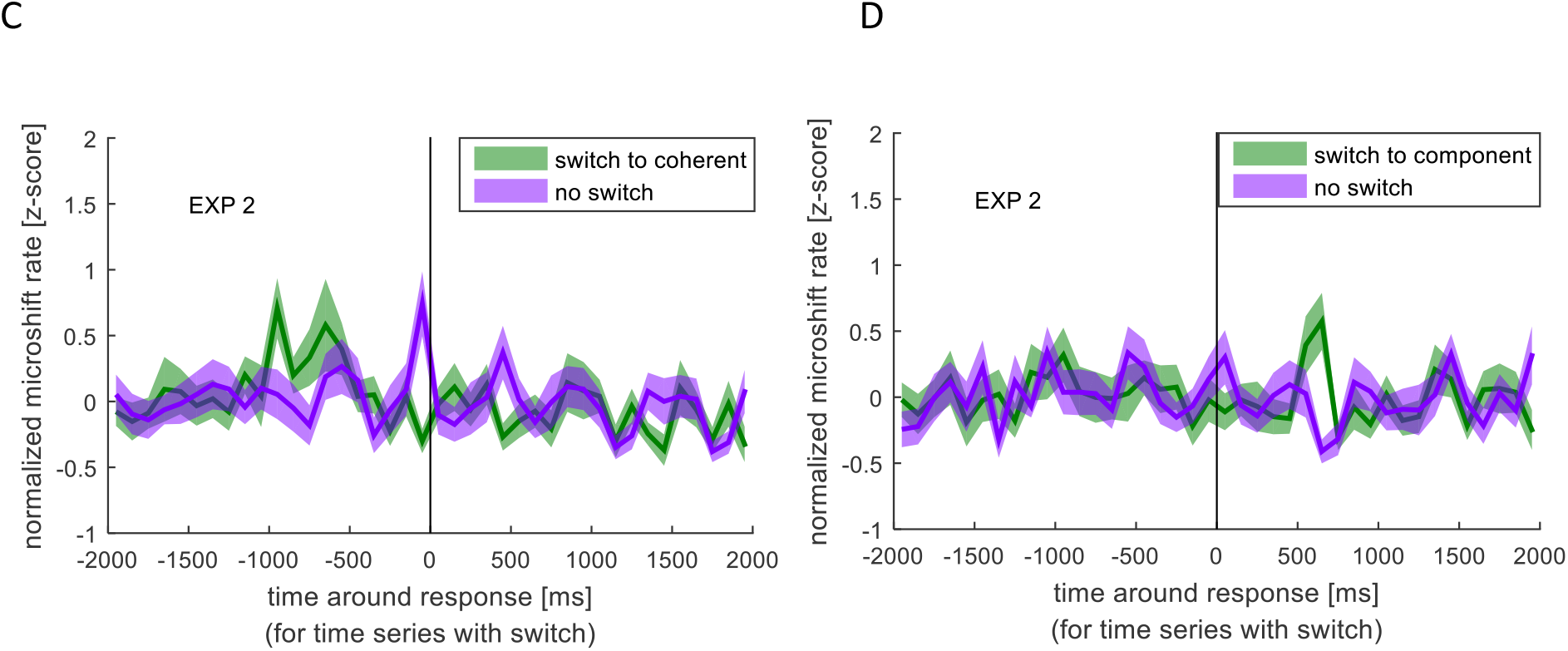
Normalized microsaccade/microshift rate around the response indicating a perceptual switch (red/green) compared to the normalized microsaccade/microshift rate during no switch (blue/ purple). Colored lines and ribbons represent mean ± SEM. Vertical shaded area marks significant time points revealed by the non-parametrical statistical test procedure described by Maris & Oostenveld (2007). A. Microsaccade rate around switch to coherent motion. B. Microsaccade rate around switch to component motion. C. Microshift rate around switch to coherent motion. D. Microshift rate around switch to component motion.

Similar to the blink rate decrease before a switch, we found a significant decrease in microsaccade rate between −300 and 0 ms before the switch to component motion. However, such a decrease was not visible before a switch to coherent motion. Microshifts showed a different pattern, which resembles the blank rate pattern in respect to the increase before a switch to coherent motion, but the difference between normalized microshift rate before or after any switch compared to no switch was not significant.

### Microsaccade direction

During the perception of the stimulus, fast eye movements with typical microsaccadic characteristics could be observed in the direction opposite to the stimulus motion. Independent of the percept, we found that the main direction of microsaccades was opposite to the coherent motion direction. After calculating the percentages of microsaccades for all directions in steps of 10° (Fig. 6), we found 21.99 % of all microsaccades during coherent percept directed opposite to the physical movement of the coherent motion (180° ± 10°), but also 15.53 % during component percept share this direction.

**Figure 6.**
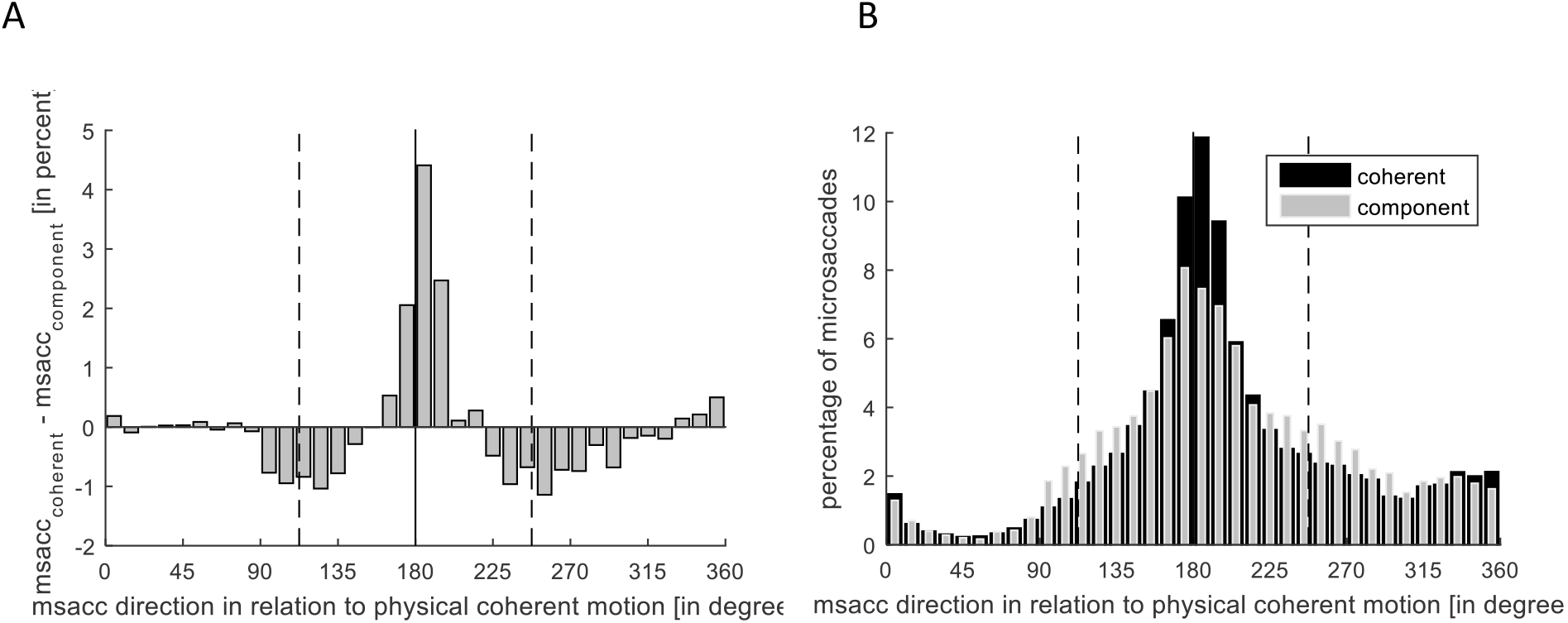
A. Microsaccade direction presented as the difference in microsaccade number during coherent motion and component motion perception in percent. B. Microsaccade direction presented as microsaccade number in percent during coherent (black) and component motion (grey) perception, separately. The solid line marks the direction of the coherent motion, while dashed lines mark the directions of component motion.

Despite this clear dominance of the direction opposite to the coherent motion, there was an influence of the percept on the distribution of the microsaccade direction circumstantiated by the significantly higher percentage of microsaccades in this direction during coherent percept than during component percept (*t*(18) = 3.57, *p* = .002, *d* = 0.82).

Accordingly, significantly more microsaccades were directed opposite to the component motions (112.5°±10° and 247.5°±10°) when component motion was perceived compared to when coherent motion was perceived (*t*(18) = −5.84, *p* < .001, *d* = 1.34). On average, 20.26 % of all microsaccades directions differed between the percepts.

### Time resolved microsaccade rate around blinks, blanks and microshifts

Normalized microsaccade rate (within 50 ms bins) around external sensory changes (blanks and microshifts) and internally introduced sensory changes (blinks) is depicted in figure 7 aligned to either blink or blank on- or offset. Microshifts consisted of a change between two frames, so the onset is equal to the offset. A pronounced reduction in microsaccade rate could be observed around all events. However, the microsaccade rate decrease started at different time points for external events (blank and microshift) compared to the internally introduced blinks. While the rate dropped immediately after the onset of the (unpredictable) blanks and microshifts, the decrease started already 200 ms before a blink. The quick closure and opening of the eye during a blinks can lead to the wrong detection of saccadic events. However, due to the observed long alteration in microsaccade rate around blinks, our conservative exclusion of microsaccades 20 ms around a blink (see methods) is unlikely to have influenced this outcome.

**Figure 7.**
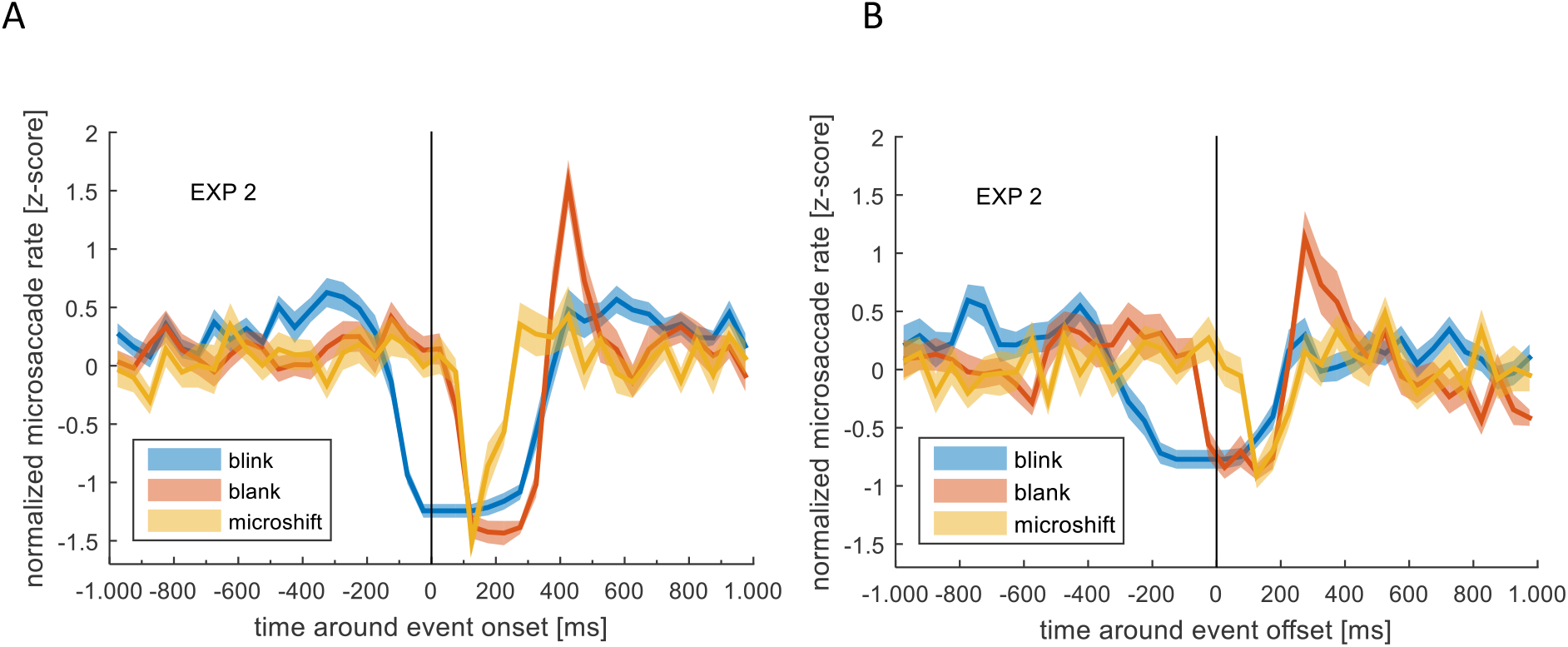
A. Normalized microsaccade rate around blink **onset** (blue), blank **onset** (red) and microshift (yellow). B. Normalized microsaccade rate around blink **offset** (blue), blank **offset** (red) and microshift (yellow). Colored lines and ribbons represent mean ± SEM.

Looking at the time after the onset of events, there was a clear peak in microsaccade rate at approx. 400 ms after blank onset, but not for microshifts or blinks. Note that the reduction before blink onset is a real rate modulation, while the low rate after blink onset is due to the fact that the eye is closed and therefore no microsaccades can be detected using a video-based eye tracker.

## Discussion

We examined the time-resolved frequency of eye blinks and microsaccades during perceptual bistability. We found that eye blinks decrease before and increase after the reported perceptual switch. When examining the two types of perceptual switches (coherent vs component motion) separately, there was a difference in modulation, with only the switch to coherent motion being accompanied by an increase after the perceptual switch report. Microsaccades, also showed a percept specific modulation in their rate, with a decrease specifically before the report of the switch to the component percept. Additionally, microsaccade direction reflected the perceived motion direction. When mimicking the visual consequences of blinks and microsaccades by introducing a transient visual interruption (blank) or a small shift of the stimulus (microshift), we found that a blank would significantly introduce a switch of percept, however, only towards coherent motion. Interestingly, a specific interaction between blinks, blanks and microshift was found with respect to the microsaccade rate. While all events led to a significant reduction in microsaccade rate, this reduction started notably before the onset of a blink. Additionally, a subsequent increase in microsaccade rate above the baseline was found only for the blank.

### Overall rate changes

Independent of the fact that the length of the percept was significantly longer for coherent compared to component motion, we found a significantly higher overall rate of blinks and microsaccades for coherent motion. Our further results indicate that this difference might be explained by the modulation over time with respect to the switch event, as discussed below.

### Temporal modulation of blinks dependent on the perceptual event

With regard to the time resolved modulation in blinks, the manual response indicating a switch was either preceded by a decrease or followed by an increase in eye blink rates depending on the type of percept subjects switched to. Specifically, the coherent percept was accompanied with a decrease before and an increase after the switch report, whereas, the component percept only showed a decrease beforehand. However, note that the decrease before the switch report to component motion, albeit still visible, did not reach significance in the second experiment albeit higher power. This could indicate a weaker or less stable effect compared to the switch to coherent motion. Nevertheless, the missing increase in blink rate after the perceptual switch to component motion clearly depicts a percept specific modulation and suggests that blinking reflects percept specific cognitive processes.

Previous studies have reported that blinking was decreased before and increased after a response indicating a perceptual switch in a sensory bistability task (Ito et al., 2003; van Dam & van Ee, 2005, 2006). However, a specific difference with respect to the type of switch has not been addressed before. Ito et al. (2003) argue that blink modulation reflects response preparation. This has been further supported by, van Dam and van Ee (2005) who found that blinks increase, not just for the perceptual switch report, but also for random button presses. Similar results with regard to manual key presses, though in a different paradigm, were found by Cong et al. (2010), who showed that blinks are entrained by rhythmic finger tapping. The differences between the modulation of blinks around the two percepts in our study reveal that it is not a mere representation of the motor response. In the following sections, we discuss the modulation of blinks before and after the response and the percept specific and attentional effects.

#### Increase of blink rate after the indicated perceptual switch

An increase in blink rate occurred exclusively after the indicated switch to the coherent motion around the time of the button press. Many studies have found that at the end of an attentional period or at the end of task relevant information, there is an enhancement of blinking (Wascher et al., 2015). The reporting of the switch in our study could be considered the end of an attentional period or a task relevant perceptual event. However, this does not explain why only the switch to the coherent percept is associated with an increase in blink rate even though both switches are equally relevant for the task. It is therefore an interesting consideration that this difference in blinking reflects a difference in how the two percepts are induced. The two interpretations are qualitatively different, in the sense that the component motion requires an additional computation of depth, which the coherent motion does not require. Further studies are needed to understand if the increase in blinks reflects such a computational bias.

Another possible explanation for the difference between coherent and component percept could lie in a difference of perceptual dominance. It has been shown that there is a suppression of eye blinks during and following a surprise input and during perceptual transients (Bonneh et al., 2016). In our experiments, the switch to the component motion, being the less likely percept, could be such a perceptual surprise. It is therefore feasible that the perceptual dominance of coherent motion influences how much attention is drawn towards a perceptual switch and the less likely one (switch to component) leads to a suppression of blinks due to attentional allocation.

It is important to note that the exclusivity of the blink modulation for one specific percept was not due to differences in the physical direction of the perceived motion direction. We addressed this by changing the direction of the stimulus. This was done to understand 1) if the vertical movement of the eyes during a blink (Collewijn et al., 1985) are linked to the perceived motion and 2) if the direction of the two motion percepts matters. It is known that the visual system is biased towards the cardinal directions, on a neuronal level and a perceptual one (Girshick et al., 2011; Schluppeck & Engel, 2010). If the coherent motion is following a cardinal direction while the component one is not, this could cause the system to treat perceptual interpretations differently. However, we found that the direction of the perceived motion had no significant effect on the blinking pattern. Hence, the difference between the percepts is, most likely, not due to any preference of physical directions, but rather, due to some internal process.

#### Decrease of blink rate before the indicated perceptual switch

Additionally, we found a decrease in blink rate before the response indicating a switch to either percept. However, it is hard to tell if this decrease is percept specific, since it does not reach significance before the reported switch to component percept in the second experiment, albeit higher power. This could indicate a weaker or less stable effect compared to the switch to coherent motion.

In general, both perceptual changes are internal events in our task, which needed to be reported and therefore should have drawn attention towards them. Studies have reported that people tend to suppress their blinks during moments of high attention or when relevant information is being presented (Hoppe et al., 2018),even before the onset of a task-relevant stimulus (Hoppe et al., 2018; Veltman & Gaillard, 1998). This suppression occurs even for stimuli outside the visual modality (Bauer et al., 1985). This indicates the involvement of a more general, vision-independent attentional mechanism. Therefore, the allocation of attentional resources caused by the requirement to report the switch might have introduced the decrease in blink rate before the response. This interpretation however would mean that the decrease happened as a consequence of the perceptual switch.

Another possibility is that the suppression in blinking is not only a result of the perceptual switch but rather or also the cause. Indeed, increased fixation duration has been shown to lead to perceptual switching in other studies (Ellis & Stark, 1978) and since blinking interrupts fixation, the suppression of blinks might introduce a perceptual switch. Unfortunately, it is difficult to conclude with certainty as to which event, the switch or the reduced blinking, occurred first, simply because there was no objective measure of the internal perceptual switch itself. However, we have a strong indication as to when the perceptual switch happened by looking at the blank results (Fig. 4). Here, it is clear that the blank must have introduced the switch and not the other way around. Figure 3 clearly shows that the time of blink reduction before the perceptual switch overlaps with the time when the switch-introducing blank occurred. This could suggest that the perceptual switch is a consequence of the blink reduction rather than the cause.

### Modulation of microsaccades (rate and direction)

In experiment 2, we looked at the role of microsaccades in the ambiguous plaid stimulus and controlled for their retinal consequence by adding microshifts to the stimulus. We found that the overall microsaccade rate was higher for the coherent than the component percept. Additionally, we found a reduction in microsaccade rates specifically before the switch to the component percept. Please note that the discrepancy to other studies, reporting that an increase in microsaccades can introduce a perceptual switch, is most likely due to the different ambiguous stimuli used (Otero-Millan et al., 2012; Troncoso et al., 2008). These studies used the rotating snakes and the Enigma illusion, both of which alternate between movement and stationary percepts. Hence, it is likely that microsaccades specifically facilitate a switch to the motion percept. The plaid stimulus, as used by us, does not have switches between movement and no movement percept but switches between different types of motion. Using a more comparable ambiguous apparent motion stimulus a reduction before a perceptual switch has been reported before (Laubrock et al., 2008). These authors further argued that the microsaccade modulation might possibly precede the internal switch, indicating a possible causal role of microsaccades. As discussed above, we believe that our observed blank introduced perceptual switch is a strong indication as to when the perceptual switch happened with respect to the response, namely between −900 and −500 ms before the response (Fig. 4). The timing of reduction in microsaccade rate as shown in figure 5 (between −300 and 0 ms) therefore suggests that the effect happened between the perceptual switch and the response, given an average reaction time of about 0.5s to 0.7s to an actual external stimulus change (Baker & Graf, 2010; Laubrock et al., 2008; van Dam & van Ee, 2005, 2006). Although it is not possible to tell with absolute certainty if the decrease in microsaccade rate follows the internal switch, the decrease only before the response indicating a switch to component percept argues against a mere consequence of response preparation.

Interestingly, we found that the direction of microsaccades was influenced by the ongoing percept. Specifically, as shown in figure 6, we found that while the overall direction was mainly opposite to the coherent motion, this number was reduced during component percept and at the same time, the number of microsaccades in the direction opposite to the two possible component motion directions was increased. Since the direction is opposite to the percept, it is likely that the percept draws the eyes in the direction of perceived motion and the detected microsaccade is a saccade back to the required position of fixation. This could mean that the microsaccades we observe are some sort of small optokinetic nystagmus. (OKN) which is a well-known phenomenon that is triggered by moving background stimuli introducing optic flow. It consists of a slow phase in the direction of the optic flow and a short, fast jump back towards the center of the visual field. OKN is greatly reduced if visual fixation is demanded (Murphy et al., 1975), however, even during fixation of a stationary target, small eye movements, affected by a moving background, can be observed. For instance, Re et al. (2019) found that microsaccade directions are influenced by and correspond to the direction of moving dot clouds that are attended during fixation. While Laubrock calls them “OKN-like rudiments” (Laubrock et al., 2008), Pola and colleagues note that these residual movements have a rather complex relationship with the OKN (Pola et al., 1995). Further studies will need to clarify if the direction of microsaccades are a consequence of the percept or lead to the specific perceptual interpretation. What, however, is clear from our results is that the microsaccade direction and the perceptual interpretation of sensory input are not independent from each other.

### The effect of external events: blanks and microshifts

Blanks and microshifts are external events that were used as controls for the visual consequences of blinks and microsaccades. Interestingly, they have a different effect on perceptual bistability compared to their corresponding eye movements. One main finding was that the blanks introduced a switch to the coherent motion in experiment 2. Please note that the effect is also clearly visible in experiment 1 (Fig. 4), but the lower power in experiment 1 might have prevented significance. Two questions arise through the finding: 1. Why do blanks but not blinks introduce a switch despite their similar visual consequences and 2. Why do blanks specifically introduce a switch to the coherent percept?

With regard to the first question, one should bear in mind that even though blinks and blanks have a similar consequence on the retinal image, they are intrinsically different (Deubel et al., 2004; Golan et al., 2018; Higgins et al., 2009). Deubel et al. (2004) found that adding a blank after a saccade can counteract the reduced detection of target displacement due to saccadic suppression, but a blink after a saccade does not have the same effect. A similar finding was also reported for blink suppression, wherein introducing a blank period after a blink reduces the displacement suppression. The idea is that an external interruption due to a blank introduces a need to recompute the post-saccadic target location; whereas, if the interruption is due to a blink, no such need is generated (Higgins et al., 2009). In other words, interruptions or small changes during blinks are generally ignored (Maus et al., 2017). This means that the oculomotor system treats an internal event such as a blink, differently from a blank. A difference between the two is also found on a neural level. A higher activity in several visual areas have been reported for blanks and not blinks (Gawne & Martin, 2000, 2002; Golan et al., 2018) and blinks (both voluntary and spontaneous) along with self-initiated blanks are associated with a decrease in activity in higher visual areas; whereas unpredictable external darkening cause an increase in higher level areas (Golan et al., 2018). A difference in perceptual consequence following a blink vs a blank is therefore not surprising.

However, the specificity of the perceptual change due to a blank, is somewhat surprising. Blanks often led to a switch to coherent percept. This was the preferred interpretation of the stimulus. This could indicate that if a certain interpretation of a sensory input is preferred, anything that causes one to reassess/ recompute the input will tend to switch the perceptual interpretation to the preferred one. Once we reach this preferred perceptual interpretation we are more likely to blink, assuming that all relevant information has been assessed. It is important to point out that our findings might be specific for the ambiguous plaid stimulus were the two percepts are clearly based on different internal processes. While the component percept interprets the stripes separately due to different depth, the coherent percept is based on an integration over the two stripe stimuli. The investigation of other bistable stimuli can clarify this specificity.

With regard to microshifts, we did not find a significant modulation. However, a small increase before the switch to coherent motion is visible (Fig. 5D), which could indicate that they indeed facilitate specific perceptual switches comparable to blanks.

### Modulation of microsaccade rate around internal and external events

We found that the microsaccade rate, albeit mostly constant around perceptual switches, was modulated around blanks, microshifts and blinks, but with a difference in temporal dynamics around the internal (blinks) versus the external (microshifts and blanks) events. Specifically, though there was a microsaccade reduction for approximately 250 ms after the event offset, this decrease started only after the onset of external events, but clearly before the onset of the internal event.

With regard to the blanks, the modulatory pattern introduced by the blank followed the typical microsaccade rate signature, characterized by a decrease, followed by an increase and a return to baseline (Bonneh et al., 2016). This modulation has been observed during other tasks and was suggested to be the reaction to sudden changes in visual input, such as display changes as well as to internal attention capturing processes (Betta & Turatto, 2006; Engbert & Kliegl, 2003; Gao et al., 2015; Pastukhov et al., 2013). Our blanks interrupted the visual information intake likely leading to a reassessment of visual input, which required the allocation of attention. However, microshifts and blinks did not show an increase in microsaccade rate after the decrease. A possible explanation stems from the fact that during the shift, there still is visual input, whereas during the blank, there is no visual information at all. This might generate a stronger need for reevaluation during the latter. This again would not happen after a blink, since a blink is self-introduced and provides no reason to assume that the input has changed. With regard to the internal blink event, we found that the actual decrease started earlier than for the external events, namely around 200 ms before blink onset. It has been shown that microsaccades are suppressed when there is an expected visual stimulus followed by a response (Betta & Turatto, 2006). An expected change in sensory input due to a blink could trigger the same mechanism. However, it must be noted that the suppression reported by Betta and Turatto (2006) was specific to sensory information that should trigger a motor response and therefore linked to response preparation, as argued by authors. The predicted sensory change caused by a blink is not task relevant but, as has been established in the literature, is ignored by the system. That no microsaccades were detected during a blink is a result of our video-based eye tracker, where it is not possible to detect microsaccades during eyes closed. We conclude that, although the reason for the suppression of microsaccades before blink onset is not completely clear, our findings clearly indicate an interaction between blinks and microsaccades.

### Summary and Conclusions

Our study on blinks and microsaccades during a visual bistable task indicates that the execution of these eye related movements is related to internal perceptual processes, and that they additionally depend on each other. The fact that different perceptual interpretations of the same sensory input are accompanied by a specific eye movement pattern further suggests that the analysis of eye movements can differentiate between distinct cognitive processes that might otherwise go undetected.

## References

Adelson, E. H., & Movshon, J. A. (1982). Phenomenal coherence of moving visual patterns. Nature, 523–525.

Baker, D. H., & Graf, E. W. G. W. (2010). Extrinsic factors in the perception of bistable motion stimuli. Vision Research, 1257–1265.

Bauer, L. O., Strock, B. D., Goldstein, R., Stern, J. A., & Walrath, L. C. (1985). Auditory Discrimination and the Eye Blink. Psychophysiology, 636–641.

Betta, E., & Turatto, M. (2006). Are you ready? I can tell by looking at your microsaccades. Neuroreport, 1001–1004.

Binder, M. D., Hirokawa, N., & Windhorst, U. (2008). Aperture Problem. In Encyclopedia of Neuroscience. Springer.

Bonneh, Y. S., Adini, Y., & Polat, U. (2016). Contrast sensitivity revealed by spontaneous eyeblinks: Evidence for a common mechanism of oculomotor inhibition. Journal of Vision, 16(7), 1–1.

Brainard, D. H. (1997). The Psychophysics Toolbox. Spatial Vision, 433–436.

Collewijn, H., van der Steen, J., & Steinman, R. M. (1985). Human eye movements associated with blinks and prolonged eyelid closure. Journal of neurophysiology 11–27.

Cong, D. K., Sharikadze, M., Staude, G., Deubel, H., & Wolf, W. (2010). Spontaneous eye blinks are entrained by finger tapping. Hum Movem Sci, 1–18.

Deubel, H., Bridgeman, B., & Schneider, W. X. (2004). Different effects of eyelid blinks and target blanking on saccadic suppression of displacement. Perception & Psychophysics, 772–778.

Einhauser, W., Stout, J., Koch, C., & Carter, O. (2008). Pupil dilation reflects perceptual selection and predicts subsequent stability in perceptual rivalry. PNAS, 1704–1709.

Ellis, S. R., & Stark, L. (1978). Eye movements during the viewing of Necker cubes. Perception, 575–581.

Engbert, R., & Kliegl, R. (2003). Microsaccades uncover the orientation of covert attention. Vision Research 1035–1045.

Gao, X., Yan, H., & Sun, H.-j. (2015). Modulation of microsaccade rate by task difficulty revealed through between- and within-trial comparisons. Journal of Vision, 1–15.

Gawne, T. J., & Martin, J. M. (2000). Activity of primate V1 cortical neurons during blinks. Journal of neurophysiology, 84(5), 2691–2694.

Gawne, T. J., & Martin, J. M. (2002). Responses of primate visual cortical neurons to stimuli presented by flash, saccade, blink, and external darkening. Journal of neurophysiology, 88(5), 2178–2186.

Girshick, A. R., Landy, M. S., & Simoncelli, E. P. (2011). Cardinal rules: visual orientation perception reflects knowledge of environmental statistics. Nature Neuroscience 926–932.

Golan, T., Grossman, S., Deouell, L. Y., & Malach, R. (2018). Widespread suppression of high-order visual cortex during blinks and external predictable visual interruptions. bioRxiv

Higgins, S. J., Irwin, D. E., Wang, R. F., & Thomas, L. E. (2009). Visual direction constancy across eyeblinks. Attention, Perception, & Psychophysics, 1607–1617.

Hoppe, D., Helfmann, S., & Rothkopf, C. A. (2018). Humans quickly learn to blink strategically in response. PNAS, 2246–2251.

Hupé, J M, & Rubin, N. (2004). The oblique plaid effect. Vision Research, 489:500.

Ito, J., Nikolaev, A. R., Luman, M., Aukes, M. F., Nakatani, C., & van Leeuwen, C. (2003). Perceptual switching, eye movements, and the bus paradox. Perception, 681–698.

Kleiner, M., Brainard, D., & Pelli, D. (2007). “What’s new in Psychtoolbox-3”. Perception, 1–16.

Laubrock, J., Engbert, R., & Kliegl, R. (2005). Microsaccade dynamics during covert attention. Vision Research, 45(6), 721–730.

Laubrock, J., Engbert, R., & Kliegl, R. (2008). Fixational eye movements predict the perceived direction of ambiguous apparent motion. Journal of Vision 1–17.

Leopold, D. A., Wilke, M., Maier, A., & Logothetis, N. K. (2002). Stable perception of visually ambiguous patterns. Nature neuroscience, 5(6), 605–609.

Leys, C., Ley, C., Klein, O., Bernard, P., & Licata, L. (2013). Detecting outliers: Do not use standard deviation around the mean, use absolute deviation around the median. Journal of Experimental Social Psychology, 764–766.

Maris, E., & Oostenveld, R. (2007). Nonparametric statistical testing of EEG-and MEG-data. Journal of neuroscience methods, 164(1), 177–190.

Martinez-Conde, S., Macknik, S. L., Troncoso, X. G., & Dyar, T. A. (2006). Microsaccades Counteract Visual Fading during Fixation. Neuron 297–305.

Maus, G. W., Duyck, M., Lisi, M., Collins, T., Whitney, D., & Cavanagh, P. (2017). Target displacements during eye blinks trigger automatic recalibration of gaze direction. Current Biology, 27(3), 445–450.

McCamy, M. B., Collins, N., Otero-Millan, J., Al-Kalbani, M., Macknik, S. L., Coakley, D., Troncoso, X. G., Boyle, G., Narayanan, V., & Wolf, T. R. (2013). Simultaneous recordings of ocular microtremor and microsaccades with a piezoelectric sensor and a video-oculography system. PeerJ, 1, e14.

Murphy, B. J., Kowler, E., & Steinman, R. M. (1975). Slow oculomotor control in the presence of moving backgrounds. Vision Research, 15(11), 1263–1268.

Nakatani, H., Orlandi, N., & van Leeuwen, C. (2011). Precisely timed oculomotor and parietal EEG activity in perceptual switching. Cogn Neurodyn, 399–409.

Nakatani, H., & van Leeuwen, C. (2005). Individual differences in perceptual switching rates; the role of occipital alpha and frontal theta band activity. Biol Cybern, 343–354.

Necker, L. A. (1832). Observations on some remarkable optical phaenomena seen in Switzerland; and on an optical phaenomenon which occurs on viewing a figure of a crystal or geometrical solid. Philosophical Magazine and Journal of Science, 329–337.

Noest, A., Van Ee, R., Nijs, M., & Van Wezel, R. (2007). Percept-choice sequences driven by interrupted ambiguous stimuli: a low-level neural model. Journal of Vision, 7(8), 10–10.

Otero-Millan, J., Macknik, S. L., & Martinez-Conde, S. (2012). Microsaccades and Blinks Trigger Illusory Rotation in the “Rotating Snakes” Illusion. The Journal of Neuroscience, 6043–6051.

Pastukhov, A., Vonau, V., Stonkute, S., & Braun, J. (2013). Spatial and temporal attention revealed by microsaccades. Vision Research 45–57.

Pelli, D. D. (1997). The VideoToolbox software for visual psychophysics: Transforming numbers into movies. Spatial Vision, 437–442.

Pola, J., Wyatt, H. J., & Lustgarten, M. (1995). Visual fixation of a target and suppression of optokinetic nystagmus: effects of varying target feedback. Vision Research, 1079–1087.

Re, D., Inbar, M., Richter, C. G., & Landau, A. N. (2019). Feature-Based Attention Samples Stimuli Rhythmically. Current Biology 693–699.

Rubin, E. (1921). Visuell Wahrgenommene Figuren. Gyldendalske Boghandel.

Schluppeck, D., & Engel, S. A. (2010). Oblique effect in human MT+ follows pattern rather than component motion. Journal of Vision, 282–282.

Troncoso, X. G., Macknik, S. L., Otero-Millan, J., & Martinez-Conde, S. (2008). Microsaccades drive illusory motion in the Microsaccades drive illusory motion in the. PNAS, 16033–16038.

Valsecchi, M., Betta, E., & Turatto, M. (2007). Visual oddballs induce prolonged microsaccadic inhibition. Experimental Brain Research, 177(2), 196–208.

van Dam, L. C. J., & van Ee, R. (2005). The role of (micro)saccades and blinks in perceptual bi-stability from slant rivalry. Vision Research, 2417–2435.

van Dam, L. C. J., & van Ee, R. (2006). The role of saccades in exerting voluntary control in perceptual and binocular rivalry. Vision Research, 787–799.

Veltman, J. A., & Gaillard, A. W. K. (1998). Physiological workload reactions to increasing levels of task. Ergonomics 656–669.

von Schiller, P. (1933). Stoboskopische Alternativbewegungen. Psychologische Forschung 17:179-214, 179–214.

Wallach, H. (1935). Über visuell wahrgenommene Bewegungsrichtung. Psychologische Forschung, 325–380.

Wascher, E., Heppner, H., Möckel, T., Kobald, S. O., & Getzmann, S. (2015). Eye-blinks in choice response tasks uncover hidden aspects of information processing. EXCLI journal, 1207–1218.

Zuber, B. L., Stark, L., & Cook, G. (1965). Microsaccades and the velocity–amplitude relationship for saccadic eye movements. Science, 1459–1460.

